# Discovery of cell-active small molecule inhibitors of UDP-galactose 4’-epimerase

**DOI:** 10.64898/2026.06.17.733026

**Authors:** Sai Kwan Khal, Noah A. Linhart, Shivam Jain, Gilbert Rosario Acevedo, Michael Boyce

**Author notes:** Equal contributions (Sai Kwan Khal, Noah A. Linhart). Equal contributions (Shivam Jain, Gilbert Rosario Acevedo).

## Abstract

Glycosylation depends on tightly regulated pools of nucleotide-sugars (NS), yet the mechanisms controlling mammalian NS homeostasis and their downstream effects on glycoprotein biosynthesis remain poorly understood. UDP-galactose 4′-epimerase (GALE) catalyzes the reversible interconversion of UDP-galactose/UDP-glucose and UDP-*N*-acetylgalactosamine/UDP-*N*-acetylglucosamine, making it a central regulator of glycan precursor pools and an excellent model enzyme for studying NS metabolism. Here, we report the discovery of a cell-active small molecule inhibitor of human GALE through a high-throughput chemical screening strategy. Using a coupled luminescence-based assay, we identified the FDA-approved drug disulfiram as a GALE inhibitor. Biochemical analyses demonstrated that disulfiram directly inhibits GALE through covalent modification of cysteine residues, including C153, likely via its reactive metabolite diethyldithiocarbamate. In cultured human cells, disulfiram treatment phenocopied genetic GALE deletion, reducing terminally sialylated glycans, mucin-type O-glycans, and properly glycosylated mucin-domain glycoproteins. These effects were rescued by galactose supplementation, consistent with a mechanism of on-target GALE inhibition. Similar phenotypes were observed in human lung adenocarcinoma cells, supporting a broader role for GALE in regulating glycosylation and mucin biosynthesis across tissue types. Together, these studies establish a platform for the discovery of pharmacological GALE inhibitors as new research tools, identify disulfiram as a cell-active chemical probe for studying NS regulation, and suggest that targeting GALE might modulate mucin hypersecretion in muco-obstructive diseases and mucinous cancers.

## Introduction

Glycosylation – the enzymatic attachment of carbohydrates to proteins, lipids, and other biomolecules – is one of the most abundant post-synthetic modifications in nature^1–3^. Carbohydrate moieties affect both the structures and functions of glycoproteins in a vast range of contexts and processes^4–7^. Nearly all glycosylation events rely on nucleotide-sugars (NSs), small metabolites that serve as monosaccharide donors for glycosyltransferases^8, 9^. In mammals, NSs are biosynthesized predominantly in the cytoplasm and are often imported into the endoplasmic reticulum (ER) and Golgi apparatus, where extensive glycosylation of secreted and cell surface proteins occurs^10, 11^. The interconnected mammalian NS biosynthetic pathways are well-characterized biochemically^12–15^, and NS levels fluctuate rapidly in response to many stimuli, such as feeding and ER stress^16, 17^, presumably to accommodate changes in glycoprotein biosynthetic demand. However, it remains incompletely understood how mammalian cells and tissues dynamically regulate NS pools and what the molecular and cellular consequences of NS dysregulation might be.

As a first step toward understanding how and why NS levels are regulated, we selected uridine diphosphate (UDP)-galactose 4’-epimerase (GALE) as a model enzyme^18^. Mammalian GALE catalyzes the epimerization of two pairs of NSs: the UDP-hexoses UDP-galactose (UDP-Gal) and UDP-glucose (UDP-Glc) and the UDP-HexNAcs UDP-*N*-acetylgalactosamine (UDP-GalNAc) and UDP-*N*-acetylglucosamine (UDP-GlcNAc)^15, 19–21^. Through these reversible epimerizations, GALE acts solely to balance preexisting pools of NSs, all of which can be biosynthesized via distinct *de novo* and salvage pathways^8, 15^. Given this role, GALE is an attractive model for studying dynamic NS regulation and downstream glycoprotein biosynthesis.

Further underlining its significance, the *GALE* gene is mutated in human disease. Partial loss-of-function *GALE* mutations cause galactosemia type III, a rare, inherited disorder that impairs the metabolism of dietary galactose^22–24^. Patients with type III galactosemia exhibit a wide variety of symptoms and degrees of severity, with heterogeneity likely caused by a spectrum of *GALE* mutations, dietary and lifestyle differences, and unknown genetic modifiers^24^. Due to this clinical variability and the rarity of type III galactosemia, it has been challenging to deduce the molecular and cellular effects of *GALE* mutations from human genetic information and patient histories^25^. Recently, it was reported that specific *GALE* mutations cause hypoglycosylation of the von Willebrand factor receptor glycoprotein Ib-α and β1 integrin on platelet surfaces, leading to thrombocytopenia^26–28^. However, our knowledge of GALE’s role in glycoprotein biosynthesis and physiology remains rudimentary.

We previously reported that CRISPR-Cas9-engineered *GALE*^-/-^ cultured human cells exhibit NS imbalances and completely lack detectable mucin-type glycans, carbohydrate chains that begin with the addition of α-GalNAc to the hydroxyl of serine or threonine^18^. Mucin-type glycans are found on a variety of secreted and membrane-bound glycoproteins, including the mucin family, the major solid component of mucus^10, 29, 30^. Mucins are so highly glycosylated that up to eighty percent of their molecular weight can be glycans^31^. This extensive O-glycosylation gives mucins and mucus their unique biophysical properties, which in turn are required for their physiological functions. For example, on the airway epithelium, mucins and mucus enable mucociliary clearance (MCC), an innate defense mechanism that protects the lungs against aspirated food, particulates, and pathogens^32, 33^. In healthy individuals, airway mucus contains ∼2% solids, primarily mucins. In muco-obstructive respiratory diseases, including asthma, chronic obstructive pulmonary disease (COPD), bronchiectasis, and cystic fibrosis, mucus solids increase to 3-9% or higher^32–35^. This change dramatically alters the biophysical properties of mucus, causing exponential increases in viscosity and elasticity, increased osmotic pressure, ciliary collapse, and inhibition of MCC. The result is a failure of mucus clearance, which triggers a cascade of downstream pathology^32, 35^. Moreover, mucin overexpression occurs in mucinous adenocarcinoma (MAC), a tumor subtype that arises in diverse tissues, including the airway^36–38^. MAC mucin hypersecretion promotes tumor metastasis, therapy resistance, and immune invasion, identifying it as an important part of the pathophysiology of these tumors^39, 40^. Despite the importance of mucins in health and disease, there are no FDA-approved drugs that reduce mucin biosynthesis to mitigate mucus hypersecretion, highlighting the need for novel therapeutic strategies.

We reasoned that small molecule inhibitors of GALE would be valuable research tools for studying the enzyme’s functions in NS metabolism and would enable proof-of-principle experiments to determine whether GALE is a potential drug target in muco-obstructive disorders. Prior studies have attempted to develop GALE inhibitors, but the identified compounds were either not effective against the human enzyme or were not cell-active^41, 42^, leaving a key knowledge gap. Here, we report a novel high-throughput screening approach to identify pharmacological inhibitors of human GALE. An initial pilot screen identified a lead compound, disulfiram, an FDA-approved drug for alcohol use disorder^43^, as a cell active GALE inhibitor. Disulfiram-treated cultured human cells displayed significantly reduced cell surface sialylated and O-glycans, including mucins, due to on-target GALE inhibition, phenocopying genetic *GALE* ablation. Our approach identifies disulfiram as a new tool for studying GALE function and provides a blueprint for discovering novel, potent, and specific GALE inhibitors for future research and translational investigation.

## Results

### A high-throughput screen identifies small molecule GALE inhibitors

To understand the role of GALE catalytic activity in NS regulation and mucin biosynthesis, we sought to identify small molecule inhibitors through a high-throughput screening campaign. First, we designed an *in vitro* GALE UDP-hexose epimerization activity assay because this reaction has been extensively studied kinetically, mechanistically, and structurally^44–46^. In this primary screening assay, GALE catalyzes the epimerization of UDP-Glc to UDP-Gal (Figure 1A). Then, β1,4-galactosyltransferase (B4GalT1) transfers Gal from UDP-Gal to GlcNAc to generate *N*-acetyllactosamine (LacNAc) and free UDP^47^ (Figure 1A). Finally, the commercial UDP-Glo reagent (Promega) generates UDP-dependent luminescence^48^ (Figure 1A). Due to kinetic differences between GALE and B4GalT1 (k_cat_ 36 s^-1^ and 3.24 s^-1^, respectively)^49, 50^, we used GALE at a 50-fold lower concentration than B4GalT1 to sensitize the assay to GALE inhibition.

**Figure 1.**
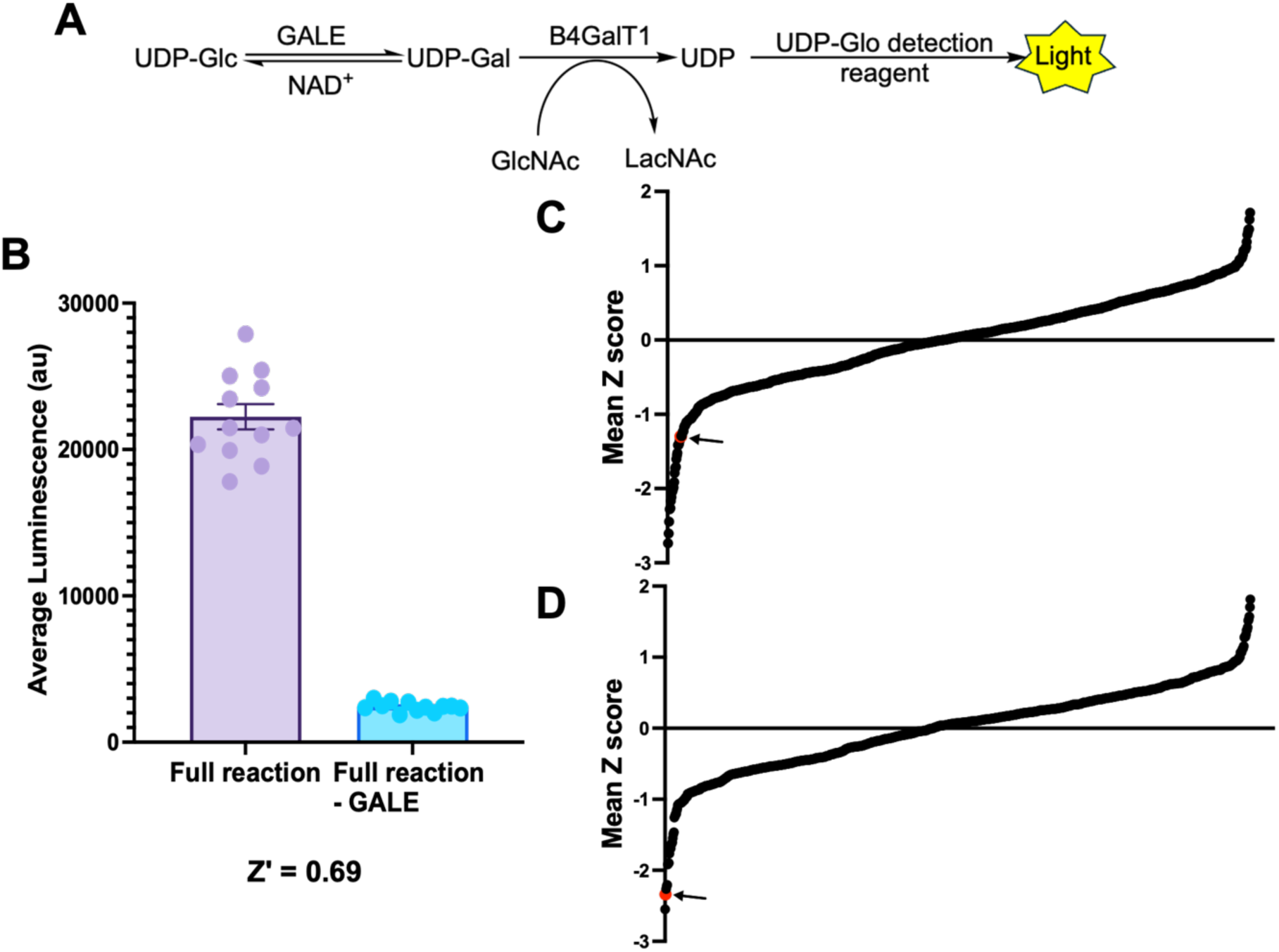
A high-throughput screening approach to identify small molecule GALE inhibitors. **A)** Primary, coupled enzymatic assay for high-throughput screening. GALE converts UDP-Glc to UDP-Gal, B4GalT1 hydrolyzes UDP-Gal, and free UDP is detected via the commercial UDP-Glo reagent and luminescence. **B)** Z-factor analysis of primary assay, comparing complete reaction and no-GALE (control) conditions. n = 3. Error bars are standard error of the mean (SEM). **C)** and **D)** Z-score data from pilot screen of 1056 compounds from an FDA-approved drug library. Compounds were screened at 100 nM **(C)** and 1 µM **(D)** in triplicate. Z-scores were calculated for each replicate and averaged. Arrows indicate disulfiram, which produced negative z-scores at both concentrations.

In initial tests, the primary assay yielded a Z’-value of 0.69, indicating that it is suitable for high-throughput screening^51, 52^ (Figure 1B). We next performed a pilot screen of the Selleck FDA-approved small molecule library (1056 compounds) in triplicate at both 100 nM and 1 µM compound concentrations. Raw luminescence values from each plate were converted to z-scores^51^ to identify potential leads. In this effort, disulfiram, an inhibitor of aldehyde dehydrogenase and an FDA-approved drug used to treat alcohol use disorder^43, 53–55^, consistently produced negative z-score values at both concentrations screened, -1.30 at 100 nM (Figure 1C) and -2.33 at 1 µM (Figure 1D). To determine whether disulfiram inhibits GALE and/or other components of the primary assay, we performed additional validation experiments, including a secondary assay without GALE and with UDP-Gal as the starting material (Figure 2A). Disulfiram exhibited dose-dependent inhibition of the primary screening assay and residual inhibition of the secondary assay but with lower potency (IC_50_ 0.902 ± 0.122 µM and 3.265 ±0.343 µM for primary and secondary assays, respectively) (Figure 2B). Taken together, these results demonstrate that our coupled *in vitro* activity assay can identify candidate inhibitors of human GALE, including disulfiram.

**Figure 2.**
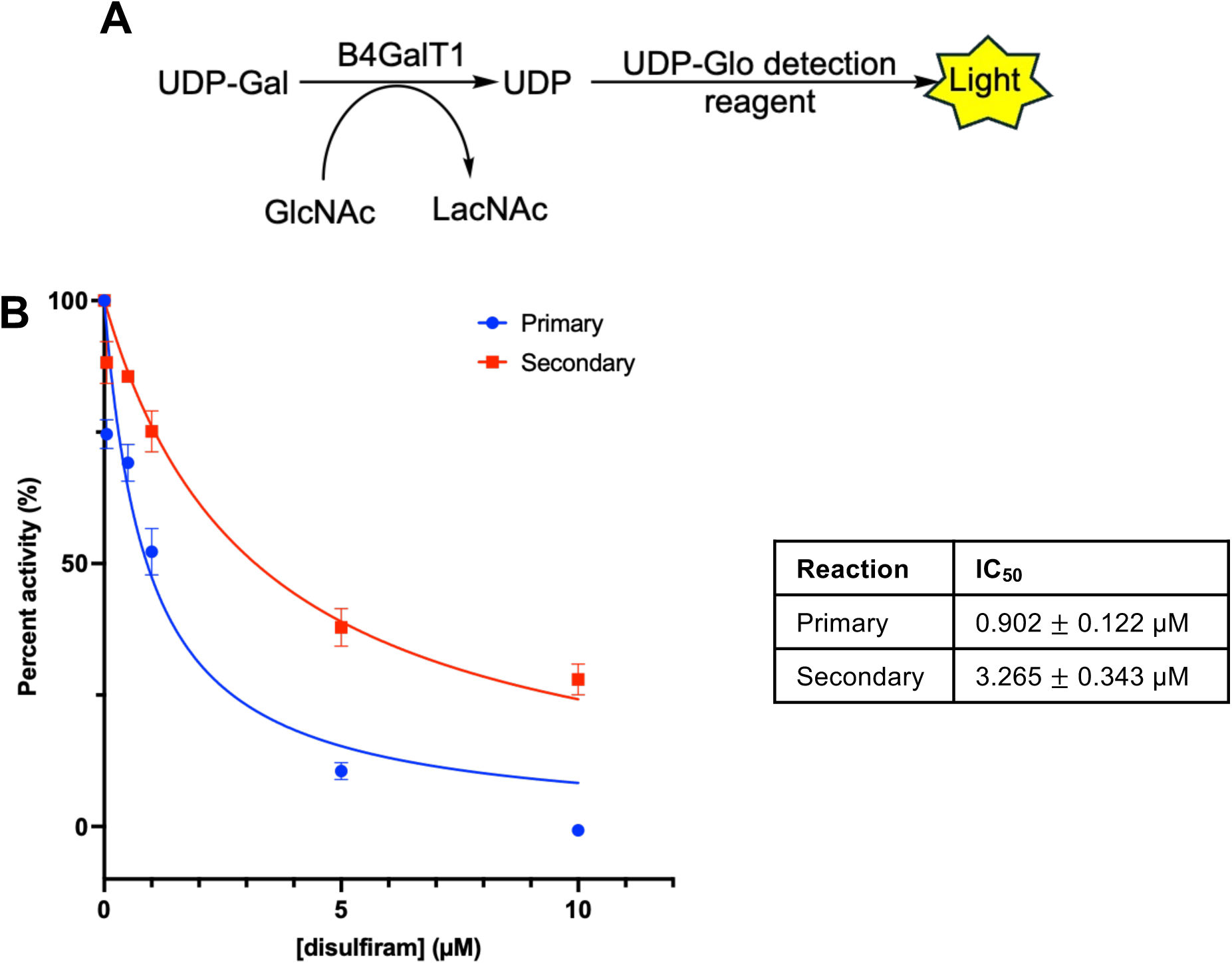
Disulfiram inhibits GALE activity *in vitro*. **A)** Secondary enzymatic assay used for validation of lead compounds. **B)** Disulfiram (0-10 µM) was tested against both primary and secondary assays to confirm that disulfiram inhibits GALE. IC_50_ values are presented as mean ± SEM. n = 3.

### Biochemical characterization of GALE inhibition by disulfiram

Disulfiram contains a reactive disulfide bond that is reportedly readily reduced by a cysteine sulfhydryl group in aldehyde dehydrogenase, yielding diethyldithiocarbamate (DDTC), which forms a covalent disulfide bond between the cysteine and DDTC, leading to enzyme inhibition^53, 55^. We hypothesized that disulfiram might inhibit GALE through a similar covalent mechanism, in which case inhibition would persist even after compound wash-out. To test this notion, we preincubated GALE with DMSO (vehicle control) or disulfiram for 30 minutes, then performed buffer exchange and activity assays (Figure 3A). We observed a significant reduction in the activity of disulfiram-treated GALE, compared to control, even after buffer exchange, consistent with a covalent inhibition mechanism.

**Figure 3.**
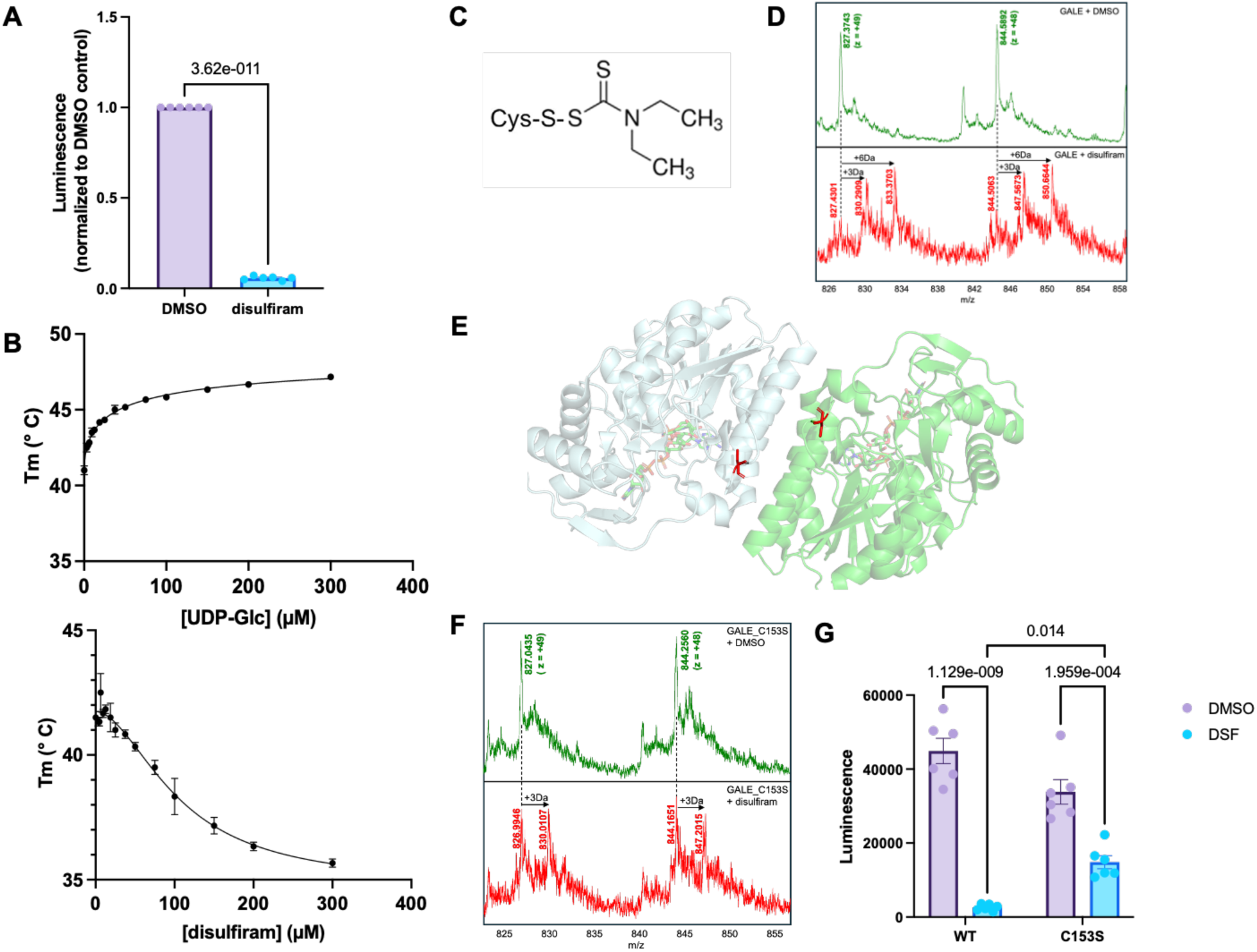
Biochemical characterization of the inhibition of GALE by disulfiram. **A)** GALE was preincubated with 200 µM disulfiram or DMSO at 30 °C for 30 min. The enzyme was then buffer-exchanged and tested for activity via the primary assay. n = 3. Data were analyzed by unpaired t-test with Welch’s correction. *p* value is provided on the graph. **B)** Thermal shift assay of GALE (10 µM) incubated with UDP-Glc (top) or disulfiram (bottom) (0-300 µM). n = 3. **C)** Addition of the disulfiram metabolite DDTC to cysteine is expected to cause a ∼147 Da mass shift. **D)** Extracted ion chromatogram at *m/z* 844.5892 ± 0.100 (*z* = + 48) and *m/z* 827.3743 ± 0.112 (*z* = + 49) of GALE (15 µM) after incubation with disulfiram (300 µM) at 30 °C for 90 minutes. n = 3, representative replicate shown. **E)** Structure of GALE homodimer (PDB 1EK6) and the location of C153 (red) adjacent to the dimer interface. **F)** Extracted ion chromatogram at *m/z* 844.5892 ± 0.100 (*z* = + 48) and *m/z* 827.3743 ± 0.112 (*z* = + 49) of GALE C153S (15 µM) incubated with disulfiram (300 µM) at 30 °C for 90 minutes. n = 3, representative replicate shown. **G)** Wild type or C153S GALE (10 µM) was incubated with DMSO or 200 µM disulfiram at 30 °C for 30 minutes and then tested in the primary screening assay. The C153S mutant is partially resistant to disulfiram-mediated inhibition. n = 3. Data were analyzed by two-way ANOVA with Tukey’s post hoc comparison. *p* values for selected comparisons are provided on the graph.

Next, we performed thermal shift assays as an orthogonal test of direct interaction between disulfiram and GALE. In this method, the binding of a small molecule results in a shift in a protein’s melting temperature (T_m_), which is reflected in the change in fluorescent intensity of the dye SYPRO Orange as it binds to exposed hydrophobic regions of protein^56, 57^. As expected, UDP-Glc, a GALE substrate, caused a dose-dependent increase in GALE T_m_ (Figure 3B). Interestingly, by contrast, disulfiram caused a dose-dependent decrease in GALE T_m_ (Figure 3B), suggesting that compound binding may destabilize the protein^58^. Because GALE is a homodimer^21, 59^, disulfiram binding may cause destabilization by promoting dimer dissociation, conformational changes, or both.

Since DDTC is reportedly the reactive metabolite of disulfiram^53^ (Figure 3C), we hypothesized that disulfiram treatment would result in a disulfide adduct between DDTC and a cysteine on GALE. To test this possibility, we examined disulfiram-treated GALE by liquid chromatography-mass spectrometry (LC-MS) (Figure 3D). LC-MS revealed the formation of two new species after disulfiram treatment, with molecular weights 147 and 294 Da larger than those in vehicle-only reactions (Figure 3D). Together, our thermal shift and LC-MS data indicate that disulfiram-derived DDTC covalently binds to GALE, likely causing or contributing to its inhibition.

Given the covalent adduct formation that we observed in disulfiram-treated GALE, we sought to identify the modified cysteine residue(s). We first tested cysteine-153 because it is surface-exposed and lies close to the dimer interface in the crystal structure of human GALE^59^ (Figure 3E). Therefore, disulfiram modification of C153 might result in conformational changes and dimer destabilization. LC-MS analysis of a C153S GALE mutant treated with disulfiram revealed the +147 Da species but not the +294 Da species observed for wild type GALE (Figure 3F), indicating that disulfiram treatment likely results in the covalent modification of C153, contributing to enzyme inhibition. Consistent with this model, the C153S mutant of GALE was partially resistant to inhibition by disulfiram, compared to the wild type enzyme (Figure 3G). In future studies, additional mutagenesis experiments and paired LC-MS and functional assays will be required to fully elucidate the mechanism of GALE inhibition by disulfiram.

### Disulfiram reduces cell-surface glycans and mucins through on-target GALE inhibition

As an FDA-approved drug, disulfiram is cell-permeable and orally bioavailable^43, 53, 54, 60^. To test whether disulfiram could be used to inhibit GALE in live human cells, we treated HeLa cells and tested for populations of glycans and mucins that are reduced in CRISPR-engineered *GALE*^-/-^ cells^18^. Vehicle- or disulfiram-treated HeLa cells were probed with *Sambucus nigra* lectin (SNA), which binds preferentially to sialic acid-containing Neu5Acα2-6Gal moieties, or jacalin, a lectin that binds preferentially to Galβ1-3GalNAc moieties, and imaged by fluorescence microscopy (Figure 4A-B). Disulfiram treatment caused a significant decrease in both SNA and jacalin staining, phenocopying genetic *GALE* deletion. To determine whether GALE is required for mucin biosynthesis, we turned to StcE-E447D, a catalytically dead mutant of the *E. coli* mucinase StcE, which is commonly employed in glycosite mapping and mucin glycoproteomics experiments because of its specificity for glycosylated mucin domains^30, 61^. StcE-E447D staining revealed that disulfiram treatment significantly reduced cell-surface mucin glycoproteins on HeLa cells (Figure 4C). From these results, we concluded that disulfiram-dependent GALE inhibition causes changes to surface glycoproteins, including mucins, in cultured human cells.

**Figure 4.**
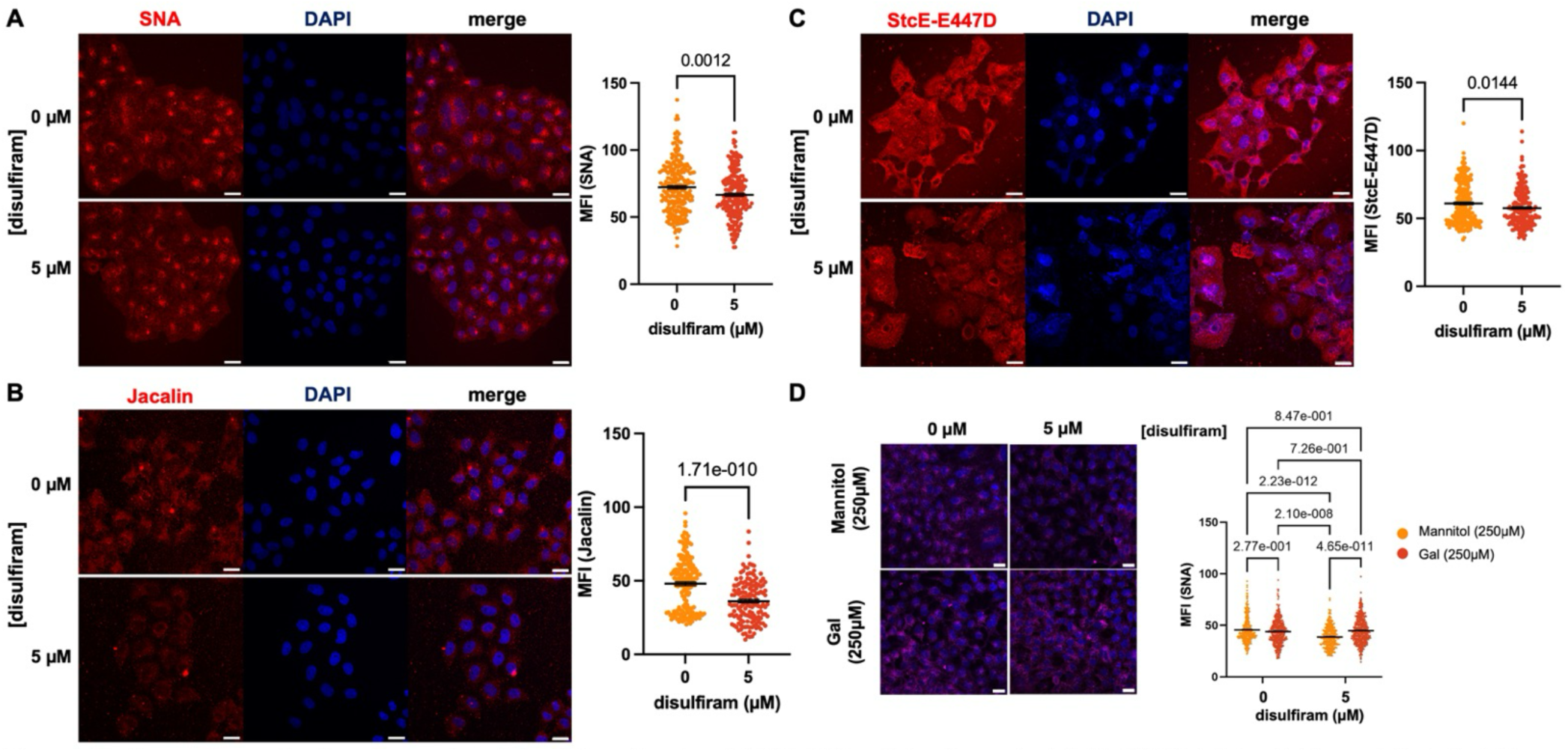
Disulfiram reduces cell-surface glycan levels through on-target GALE inhibition. HeLa cells were treated with DMSO or disulfiram as indicated for 24 hours and analyzed by fluorescence microscopy after staining with **A)** SNA or **B)** jacalin. Scale bar = 20 µm. Mean fluorescence intensity (MFI) per cell was quantified and assessed by unpaired Student’s t-test with Welch’s correction. n = 3. *p* values are provided on the graphs. **C)** HeLa cells were treated with DMSO (vehicle) or disulfiram as indicated for 48 hours and analyzed by immunofluorescence for StcE-E447D. Scale bar = 20 µm. MFI per cell was quantified and assessed by unpaired Student’s t-test with Welch’s correction. n = 3. *p* value is provided on the graph. **D)** HeLa cells cultured in 250 µM Gal or mannitol (osmolyte control) were treated with DMSO or 5 µM disulfiram for 48 hours, stained with SNA, and analyzed by fluorescence microscopy. Scale bar = 20 µm. Data were analyzed by two-way ANOVA with Tukey’s post hoc comparison. n = 3. *p* values for selected comparisons are provided on the graph.

To confirm that these effects of disulfiram treatment were caused by on-target inhibition of GALE, as opposed to off-target or nonspecific effects, we next performed rescue experiments by supplementing the medium of disulfiram-treated cells with Gal to bypass the loss of GALE function, as we have done previously with *GALE*^-/-^ cells^18^. Gal supplementation reversed disulfiram-induced changes to the glycome of HeLa cells, whereas treatment with mannitol (a non-metabolizable sugar alcohol, serving as an osmolyte control) did not (Figure 4D). We concluded that disulfiram causes a reduction in cell-surface glycan biosynthesis via GALE inhibition in HeLa cells.

### Disulfiram reduces cell-surface glycans and mucins in human lung MAC cells

Lastly, we tested whether disulfiram inhibits GALE in a tractable model of human disease. MAC is characterized by mucin hypersecretion, which promotes immune evasion and tumor growth^39, 40^. Despite this pathophysiological importance, the pathways that drive mucin hypersecretion in MAC are not fully understood. We hypothesized that GALE may be required for mucin biosynthesis in MAC. To test this hypothesis, we genetically or chemically inhibited GALE activity in human A549 cells, a manipulable model of lung MAC.

First, we generated control and *GALE*^-/-^ A549 cells using Cas9 methods and single guide RNAs (sgRNAs) targeting *GALE* or the AAVSI (“safe harbor”) locus as a control^18, 62^. Compared to controls, *GALE*^-/-^ A549 cells have drastically reduced terminally sialylated glycans at the cell surface, as judged by SNA staining, and Gal supplementation restored SNA binding in *GALE*^-/-^cells (Figure 5A). These results indicate that GALE is required for glycome biosynthesis in a MAC model. Similarly, disulfiram treatment significantly reduced SNA and StcE-E447D staining in parental A549 cells, phenocopying genetic *GALE* deletion (Figure 5B-C). These results reinforce the importance of GALE in mucin biosynthesis in diverse cell types and pave the way for future studies to determine the potential therapeutic utility of pharmacologically inhibiting GALE in lung MAC and other diseases characterized by mucin hypersecretion.

**Figure 5.**
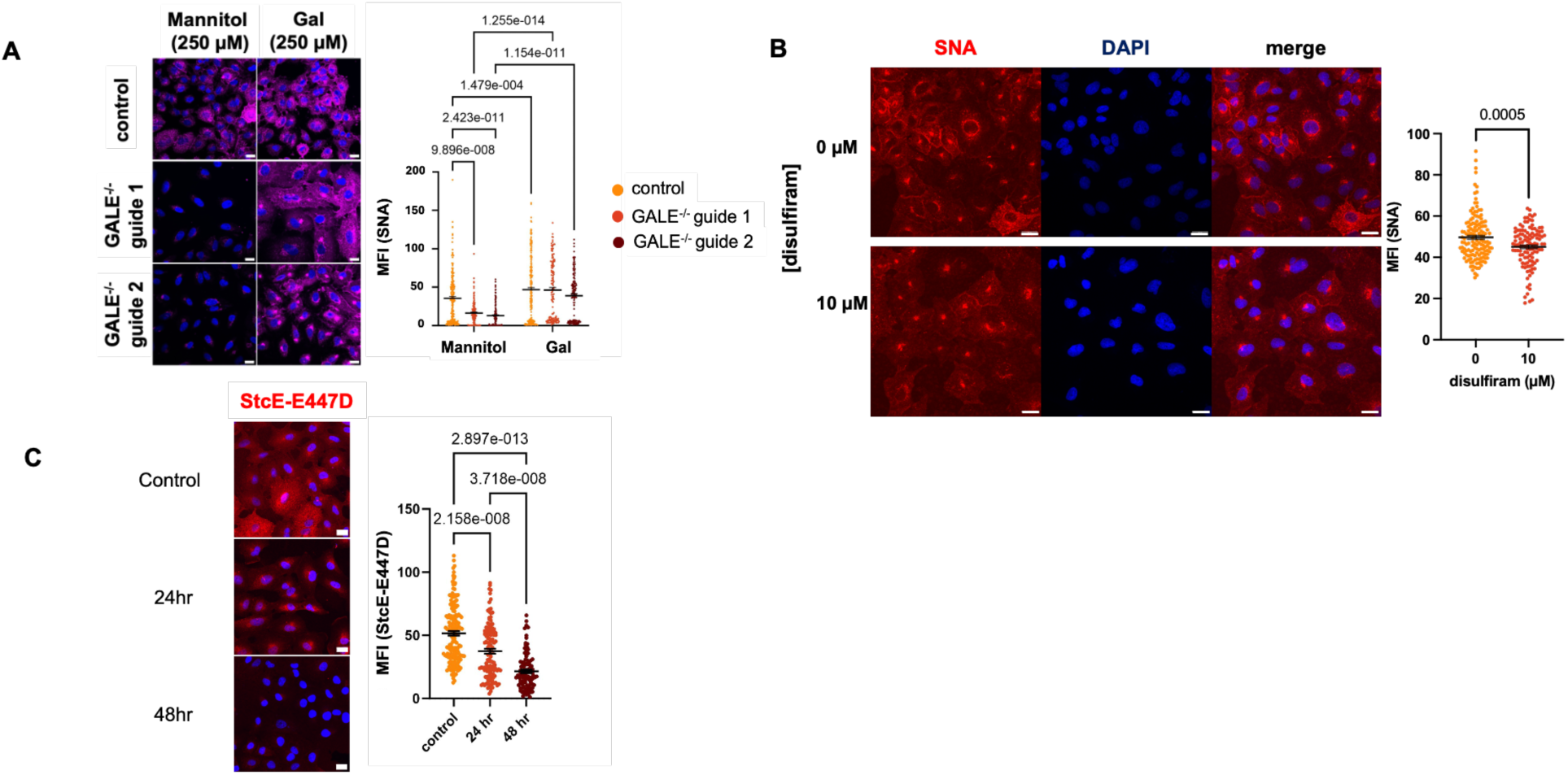
Disulfiram reduces cell-surface glycan and mucin levels in human lung adenocarcinoma cells. **A)** Control and GALE^-/-^ A549 cells were cultured in 250 µM Gal or mannitol for 24 hours and analyzed by SNA staining and fluorescence microscopy. MFI per cell was quantified and assessed by two-way ANOVA with Tukey’s post hoc comparison. n = 3. *p* values for selected comparisons are provided on the graph. **B)** Parental A549 cells were cultured with DMSO vehicle or disulfiram for 48 hours and analyzed by SNA staining and fluorescence microscopy. Scale bar = 20 µm. MFI per cell was quantified and assessed by unpaired Student’s t-test with Welch’s correction. n = 3. *p* value is provided on the graph. **C)** Parental A549 cells were cultured with DMSO vehicle or disulfiram for the indicated times and were analyzed by StcE-E447D staining and fluorescence microscopy. Scale bar = 20 µm. MFI per cell was quantified and assessed by one-way ANOVA with Tukey’s post hoc comparison. n = 3. *p* values for selected comparisons are provided on the graph.

## Discussion

Mucin glycoproteins are critical for epithelial barrier integrity, osmotic homeostasis, and protection from foreign particulates and pathogens^31, 63, 64^. However, mucin hypersecretion is implicated in various airway diseases, including asthma, allergy, COPD, bronchiectasis, cystic fibrosis, and MAC^35, 37, 63, 65–67^. Mucins owe their biophysical properties and physiological functions to their dense glycans, which are biosynthesized from tightly regulated levels of NS^64, 68^, but the metabolic pathways controlling NS pools and downstream glycoconjugate biosynthesis remain incompletely understood. Our prior work highlighted the importance of GALE for proper NS regulation and mucin glycan biosynthesis^18^. To extend these studies, we developed a high-throughput screening approach to identify chemical GALE inhibitors that will help to characterize the dynamic regulation of NSs and to investigate the potential of GALE as a therapeutic target in mucin hypersecretion diseases.

To our knowledge, disulfiram is the first reported cell-active small molecule inhibitor of human GALE. To characterize its mechanism of action, we used multiple orthogonal activity and binding assays, pointing to a covalent modification of GALE by a disulfiram metabolite, likely DDTC (Figures 3C-G). Our thermal shift data show that treatment of GALE with disulfiram leads to a decrease in the T_m_ of the protein, suggesting that disulfiram modification favors a destabilized state of GALE, perhaps promoting homodimer dissociation (Figure 3B)^58, 69^. Our LC-MS data suggest that DDTC is the active metabolite for GALE modification (Figure 3D), similar to prior reports with other targets^53–55^, and mutagenesis studies coupled with LC-MS and activity assays revealed that modification of C153 likely contributes to GALE inhibition by disulfiram (Figure 3F-G). While the detailed biophysical mechanism of inhibition awaits full characterization in future studies, we hypothesize that the covalent modification of C153 by DDTC destabilizes GALE homodimers, which contributes to the inhibition of enzymatic activity. Given our observation of two disulfiram-dependent adducts of wild type GALE in our LC-MS experiments and residual inhibition of C153S mutant GALE by disulfiram (Figures 3D, 3F-G), C153 modification does not fully explain compound action. Efforts are underway to thoroughly characterize the mechanism of inhibition of disulfiram on GALE.

Regardless of its precise mode of inhibition, our results also indicate that disulfiram can be used to manipulate GALE function in cultured human cells. For instance, in both HeLa and A549 cells, disulfiram treatment reduced cell-surface Gal-containing glycan motifs and mucins, phenocopying our prior and current results with genetically engineered *GALE*^-/-^ cells (Figures 4 and 5). In particular, StcE-E447D staining demonstrated that mucin domain-containing glycoproteins are hypoglycosylated and/or reduced in expression after 24 hours of disulfiram treatment (Figure 4C and 5C). Interestingly, in A549 cells, disulfiram treatment also appeared to cause an increase in perinuclear-localized sialylated glycans (Figure 5B), potentially reflecting the retention of hypoglycosylated proteins in the early secretory pathway. These results suggest that chemical inhibition of GALE can remodel the glycome and reduce properly glycosylated mucins relatively quickly. Moreover, the reduction in mucin expression by disulfiram was time-dependent and required re-supplementation of the compound every 24 hours (Figures 4C-D and 5C; see Materials and Methods). These observations indicate that disulfiram could be a useful tool for transiently and reversibly inhibiting GALE activity in a wide variety of experimental contexts. We note that the impact of disulfiram treatment on the cell surface glycome was more modest than that of genetic *GALE* ablation, likely because of incomplete inhibition by disulfiram, drug metabolism or efflux, and/or cellular compensation mechanisms. Optimization of disulfiram doses and treatment times will likely be required for future studies of GALE function in other systems. Taken together, these results show that loss of GALE activity causes significant changes to the glycome and reduction in mucin glycoproteins across different human cell types.

Disulfiram is a promising tool compound for studying GALE function, but it has liabilities. Perhaps most notably, disulfiram inhibition is not specific to GALE and is well-known to act on other human targets. For instance, disulfiram derives its clinical utility from inhibiting aldehyde dehydrogenase^53–55, 60^, and, in our experiments, we observed that it inhibits at least one other enzyme in our primary screening assay (Figure 2B). In the clinical setting, it may be that therapeutic doses of disulfiram achieve useful inhibition of aldehyde dehydrogenase without appreciable GALE inhibition. Importantly, in our cell-based experiments, genetically engineered *GALE*^-/-^ cells and Gal supplementation provide controls that allow us to distinguish between effects of disulfiram due to GALE inhibition versus off-target effects. It will be necessary to incorporate similar controls in all future studies of GALE function that use disulfiram. These liabilities also point to the need for more specific and potent cell-active GALE inhibitors. To this end, we have conducted a 50,000-compound screen harnessing an improved version of our *in vitro* assay, with a goal of discovering and characterizing novel GALE inhibitors in forthcoming reports.

We envision that disulfiram and other, future inhibitors of human GALE will find important uses in both basic and translational research. For example, given GALE’s importance in NS regulation and glycan biosynthesis, specific inhibitors will be powerful, titratable, and reversible tools for characterizing the mechanisms and downstream functional consequences of NS fluctuations in response to diverse stimuli, including changing nutrient availability, ER stress, and cell differentiation^16, 70–73^. Furthermore, small molecule inhibitors will enable studies testing the potential of GALE as a therapeutic target in mucin hypersecretion diseases. Mucins rely heavily on glycans for their function, and even modest changes in mucin glycoforms are predicted to have significant impacts on the viscoelastic properties of mucus. Indeed, mucus viscosity is exponentially increased by glycan-regulated mucin polymer overlap and entanglement^32, 68, 74^. Therefore, an inhaled small molecule that partially inhibits GALE activity in the airway might reduce mucin biosynthesis and mucus viscosity and provide therapeutic benefit in a wide range of mucin hypersecretion diseases, without unacceptable or toxic side-effects. Finally, beyond mucins, overexpression of cell-surface sialylated glycans contributes to cancer metastasis, immune evasion, and cell growth and proliferation^75^. Because genetic or pharmacological inhibition of GALE reduces the expression of these glycans (Figures 4A and 5B), targeting GALE might also be useful in lung MAC. In sum, disulfiram and other GALE inhibitors will enable future metabolomic, cellular, and physiological studies of how NS regulation impacts the glycome *in vivo* in both health and disease.

## Materials and Methods

### Chemicals

Disulfiram (#86720), phosphate-buffered saline (PBS; P-3813), Tris-base (T1530), GlcNAc (A8625), NAD+ (N0632), Galactose (G5388), Mannitol (M4125) and MnCl_2_ (M3634) were purchased from Sigma-Aldrich. UDP-Glc was purchased from Avantor (28053-08-9). UDP-Gal and alkaline phosphatase (M1821) were purchased from Promega (V7171).

### Cell culture

HeLa cells were grown in Dulbecco’s modified Eagle Medium (DMEM, Gibco, 11995065) containing 10% fetal bovine serum (FBS, Sigma-Aldrich, F0926), 100 units/mL penicillin, and 100 μg/mL streptomycin (Pen/Strep, Gibco, 15140122). A549 cells were gown in Ham’s F-12K (Kaighn’s) medium (Gibco, 21127022) containing 10% FBS (Sigma-Aldrich, F0926) and Pen/Strep. All cell lines were maintained at 37 °C in a humidified atmosphere with 5% CO_2_.

### GALE expression and purification

BL21 *E. coli* cells expressing codon-optimized N-terminal 6xHis-tagged human GALE in a pET-p15 expression plasmid were cultured in 10 mL LB medium with 100 µg/ml ampicillin overnight at 37 °C and 220 rpm. Then, overnight cultures were used to inoculate 1 L LB medium with 100 µg/ml ampicillin. Cultures were grown at 37 °C and 220 rpm to OD_600_ = 0.5 and then induced with isopropyl β-D-1-thiogalactopyranoside (IPTG, 0.5 mM) and grown overnight at 18 °C and 220 rpm. Bacteria were harvested by centrifugation at 3000 g for 30 min at 4 °C. The cell pellet was resuspended in BugBuster Protein Extraction Reagent (Sigma-Aldrich) and incubated for 1 hour at 4 °C for lysis. The cell lysate was centrifuged at 25,000 g for 30 min at 4 °C. The supernatant was collected, and GALE was purified using a 5 mL TALON metal affinity resin (Takara) column equilibrated with wash buffer A (20 mM sodium phosphate pH 7.4, 200 mM NaCl, 10 mM imidazole). The column was washed with 300 mL wash buffer A followed by 200 mL wash buffer B (20 mM sodium phosphate pH 7.4, 200 mM NaCl, 15 mM imidazole, 2 M urea), and 50 mL wash buffer A. GALE protein was eluted using elution buffer (20 mM sodium phosphate pH 7.4, 200 mM NaCl, 250 mM imidazole). Protein-containing fractions were identified using Bradford reagent. The elution fractions were pooled, concentrated to 3 mL, and buffer-exchanged using 10DG desalting columns (Bio-Rad) into storage buffer (20 mM sodium phosphate pH 7.4, 200 mM NaCl, 20% glycerol). The desalted protein was analyzed by SDS-PAGE and colloidal Coomassie stain, flash-frozen in liquid nitrogen, and stored at -80 °C.

### Generation of GALE C153S

C153S mutation of GALE was created in the wild type pET-p15-GALE plasmid using the following primers:

5’ CATCCGACCGGCGGTAGCACCAACCCGTATGG 3’

5’ CCATACGGGTTGGTGCTACCGCCGGTCGGATG 3’

PCR was carried out following the Phusion Hot Start II DNA Polymerase protocol (Thermo F549L), followed by DpnI digestion at 37 °C for 1 hour. The reaction was then used to transform NEB5α and BL21 (DE3) *E. coli* cells. DNA sequencing was used to verify the mutation. The mutant protein was expressed and purified as described above for wild type GALE.

### StcE-E447D expression and purification

Professor Raphael Valdivia (Duke) kindly provided the plasmid pET28a(+)-StcE-E447D. A 6x-HisFlag tag was subcloned onto the C-terminus of the protein using Gibson assembly, with primers:

5’ GACTACAAAGACGATGACGACAAGCTCGAGCACCACCACCACCACC 3’

5’ CTTGTCGTCATCGTCTTTGTAGTCTTTATATACAACCCTCATTGAC 3’.

*E. coli* BL21(DE3) were transformed with the construct and grown at 37 °C until OD 0.6-0.8. The culture was induced with IPTG (0.3 mM) and incubated overnight at 20 °C (225 rpm). Cells were collected and lysed in 20 mM HEPES pH 7.5, 500 mM NaCl using a probe tip sonicator. Lysates were centrifuged at 25,000 g for 30 min at 4 °C. Clarified cell lysates were incubated with Ni-NTA beads overnight at 4 °C. Washes were performed with increasing amounts of imidazole (10 mM to 50 mM) and 500 mM NaCl in PBS. StcE-E447D was eluted from the column with 1 mL of 250 mM imidazole and 500 mM NaCl in PBS. Pooled fractions were concentrated and buffer-exchanged using 10DG desalting columns (Bio-Rad) into storage buffer (20 mM sodium phosphate pH 7.4, 200 mM NaCl, and 20% glycerol). The desalted protein was analyzed by SDS-PAGE, flash-frozen in liquid nitrogen, and stored at -80 °C.

### High-throughput GALE small molecule screening assay

High-throughput screening assays were carried out in 384-well plates (Corning, 3574). A library of 1056 FDA-approved compounds from SelleckChem was used for pilot screening. Compounds were dissolved in DMSO and screened at final concentrations of 100 nM and 1 µM. Compounds were added to columns 1 to 22. Columns 23 and 24 were used for controls. Each plate was screened in triplicate. 25 µL of freshly prepared reagents on ice in 20 mM Tris, pH 7.5 were added to each well using a Biotek MultiFlo microplate dispenser to columns 1-23. The final composition in each well was UDP-Glc (0.1 mM), GlcNAc (2 mM), MnCl_2_ (5 mM), NAD^+^ (2 mM), GALE (1 ng), and B4GalT1 (50 ng). B4GalT1 was purchased from Glyco Expression Technologies. In columns 23 and 24, only DMSO was added. Column 23 represents negative control samples (no inhibition). In column 24, reactions without GALE served as positive controls (no signal; mimics 100% inhibition). Plates were incubated at 30 °C for 1 hour. After incubation, 7 µL of UDP-Glo detection reagent (Promega) was added to each well and incubated at room temperature (RT) for an additional 30 min before luminescence was measured using a BMG Clariostar Multimode Plate Reader.

For each plate, the Z value was calculated using the formula:

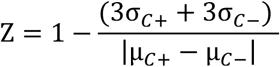

where σ*_c_*_+_ and µ*_c_*_+_ are the standard deviation and mean of the positive control samples (no GALE), and σ*_c_*_-_ and µ*_c_*_-_ are the standard deviation and mean of the negative control samples.

For columns 1-22, a z-score was calculated for each well using the formula:

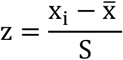

where x_i_ is the raw luminescence value in ith well, x̅ and S are the mean of luminescence and the standard deviation of all test wells without controls, respectively.

### Disulfiram wash-out experiments

GALE (10 µM) was incubated with disulfiram (200 µM) or DMSO for 30 min at 30 °C. The protein mixture was then exchanged into storage buffer using Zeba Spin Desalting Columns (Thermo 89882). Desalted GALE was then tested for activity using the primary assay as described above.

### Thermal shift assay

Thermal shift assays were performed using a CFX Connect Real-Time PCR Detection System (BioRad). Samples contained SYPRO Orange Dye (1X), GALE (10 µM), and disulfiram (0-300 µM). SYPRO Orange (400X, Thermo Scientific S6651), GALE (40 µM), and disulfiram prepared in 20% DMSO were combined in a clear 96-well PCR plate (BioRad, HSP9601) to a total volume of 10 µL. Assay was carried out between 25-95 °C. A temperature increment of 0.5 °C/min was applied, and fluorescence was measured every minute. The minimum point of the plot of the first derivative for each disulfiram concentration melt series was used to generate the T_m_ curve.

### Liquid Chromatography-Mass Spectrometry (LC-MS)

GALE (15 µM) was incubated with disulfiram (300 µM) for 1.5 hours at 30 °C. The enzyme was then buffer-exchanged into 20 mM Tris, pH 7.5, 200 mM NaCl prior to LC-MS analysis. LC-MS was performed on a 6224 TOF LC-MS system (Agilent Technologies), consisting of a 1200 HPLC (degasser, binary pump, thermostated column compartment, diode array detector) coupled to a 6224 accurate-mass time-of-flight mass spectrometer. The mass spectrometer was equipped with a Dual ESI source, and accurate mass data were obtained by internal calibration (reference ions 121.050873 and 922.009798 m/z) using a secondary nebulizer to continuously deliver the reference solution. Positive-ion mass spectral data were acquired in full-scan mode over the range of 100-3200 m/z using the following source parameters: gas temperature 325 °C, gas flow 11 L/min, nebulizer pressure 33 psig, VCap 3500 V, and fragmentor voltage 220 V. Mass spectra were acquired in centroid mode, and full scan data were processed using the deconvolution maximum entropy feature of the Agilent Masshunter Bioconfirm software. HPLC separations were achieved on a Phenomenex Kinetix EVO C18 column (3 x 100 mm, 2.6 µ) using a linear gradient of mobile phase B in A, a flow rate of 0.5 mL/min, and a column temperature of 40 °C. Mobile phase A was prepared by combining 400 mL ultrapure water with 12 mL methanol and 1.2 mL formic acid. Mobile phase B was prepared by combining 400 mL acetonitrile with 12 mL ultrapure water and 1.2 mL formic acid. The gradient program included an initial hold at 0% solvent B for 0.5 min, followed by a linear increase to 100% solvent B from 0.5-8 minutes, a hold at 100% solvent B from 8-9 minutes, and re-equilibration back to 0% B for a total run time of 15 min. Samples were analyzed using a 0.5-5-µL injection volume.

### Immunofluorescence

For 24-hour disulfiram treatment assays, 75,000 cells were seeded in 12-well plates with an 18 mm coverslip on the bottom. The following day, the medium was changed to DMEM containing disulfiram. Twenty-four hours later, cells were washed with PBS, fixed with 4% paraformaldehyde (PFA) at RT for 20 min, washed 3 times with PBS for 5 min, permeabilized with PBST (0.1% Triton X-100 in PBS) at RT for 10 min, and incubated in blocking buffer (2.5% bovine serum albumin (BSA, Sigma-Aldrich, A9647) in PBST) at RT for 30 min. Cells were washed with PBS at RT for 5 min twice. For StcE-E447D staining, cells were incubated with 20 µg/mL Flag-tagged StcE-E447D in PBS at RT for 1 hour prior to incubation with primary antibodies diluted 1:500 in blocking buffer overnight at 4 °C. The coverslips were washed with PBS at RT for 5 min thrice and then incubated with secondary antibodies conjugated with AlexaFluor dyes diluted 1:500 in blocking buffer in the dark at RT for 1 hr. Cells were washed with PBST thrice at RT for 5 min and then mounted with DAPI Fluoromount-G (SouthernBiotech 0100-20) onto microscope slides. Images were acquired on a Zeiss 780 with a 40X/1.4 NA Oil Plan-Apochromat DIC objective lens. Diode (405 nm), argon (488 nm), and helium-neon (633 nm) lasers were used for fluorophore excitations. Z-stacks were acquired for each image and processed using Fiji Image J.

For 48-hour disulfiram treatment, 37,500 cells were seeded. The following day, the medium was replaced with DMEM containing disulfiram and again 24 h later. Cells were then processed as above.

For Gal rescue experiments, 250 µM Gal or mannitol was added to DMEM 24 h before cell fixation. Cells were then processed as above.

The following primary probes were used: Anti-Flag (mouse, Sigma-Aldrich F1804); SNA (biotinylated, Vector Labs B-1305-2; Cy5 conjugated, Vector Labs CL-1305-1); jacalin (biotinylated, Vector Labs B-1155-5).

## Acknowledgements

We thank Seth Taylor and So Young Kim, Ph.D. of the Duke Functional Genomics Core for help with small molecule screening, Peter Silinski, Ph.D. of the Duke University Chemistry Department for LC-MS analysis, Dr. Kenichi Yokoyama for helpful advice, Quyen Nguyen for statistical analyses, and Brycen Aldrich and Isha Korgaonkar for technical contributions. This work was supported by grant MF-1912-00621 from the G. Harold & Leila Y. Mathers Foundation and grant R35GM161290 from the National Institutes of Health to M.B., appointment of N.A.L. to NIH training grant T32GM142605, and support from the PRIME Cancer program via NIH award R25CA275731 to G.R.A.

